# A panel of genotypically and phenotypically characterised WHO priority Gram-negative bacteria to facilitate antimicrobial and diagnostic development

**DOI:** 10.64898/2026.09.18.752562

**Authors:** Jake K Soley, Shazad Mushtaq, Charlotte Hind, Rachael Adkin, Hannah McGregor, Esther Sweeney, Ed Hutchings, Helen Mason, Elena BM Breidenstein, Lisa Dawson, Peter J Coombs, Beverley Isherwood, J. Mark Sutton, Katie L Hopkins

## Abstract

**Background:** Antimicrobial-resistant Gram-negative bacteria pose a major threat to global health. In the WHO Bacterial Priority Pathogen List (BPPL) 2024, several species were classified as ‘critical’ or ‘high’ priority for research and development of new antimicrobial therapeutics and diagnostics. However, access to well-characterised and clinically relevant bacterial isolate panels remains an unmet need for development and validation.

**Methods:** We constructed five panels of Gram-negative bacterial isolates, cultured from samples originally referred to the UK Health Security Agency Antimicrobial Resistance and Healthcare-Associated Infections Reference Unit for analysis between 2014 and 2025. For each isolate, antimicrobial susceptibilities were determined against a range of clinically relevant antibiotics, including third-generation cephalosporins, carbapenems, colistin and other ‘last-line’ antibiotics. WGS was performed on all isolates, and previously described resistance determinants were characterised.

**Results:** The five panels consist of 145 isolates and represent 81 sequence types across *Escherichia coli* (30 isolates), *Klebsiella pneumoniae* (30 isolates), carbapenem-resistant *Acinetobacter baumannii* (30 isolates), carbapenem-resistant *Pseudomonas aeruginosa* (30 isolates), and a mixed panel of other healthcare-associated Enterobacterales species (25 isolates). Panels are highly representative of globally relevant strains, including those currently in circulation and on the WHO BPPL 2024, and of resistance mechanisms of public health importance.

**Conclusion:** The panels provide a diverse, comprehensive set of multidrug-resistant Gram-negative bacterial isolates, representative of strains currently in circulation globally. Selected isolates carry a breadth of important resistance mechanisms and are highly relevant to the current epidemiological landscape. The isolates and associated metadata are available to industry, academia, and other laboratories for use in developing novel antimicrobial compounds and diagnostic assays.

## INTRODUCTION

Infections caused by Gram-negative bacteria pose a significant global healthcare challenge, exacerbated by the emergence and dissemination of antimicrobial resistance (AMR). Resistance to beta-lactam antibiotics, including cephalosporins and carbapenems, has risen in recent years and is a particular concern given their widespread use in the treatment of Gram-negative infections: deaths associated with carbapenem-resistant Gram-negative bacteria rose from 619,000 in 1990 to 1.03 million in 2021.^1,2^ Yet despite increasing rates of resistance, the rate of antibiotic discovery has slowed in recent years, particularly for agents active against Gram-negative species.^3,4^

Several antibiotic-resistant species were classified as ‘critical’ or ‘high’ priority for research and development in the WHO Bacterial Priority Pathogen List (BPPL) 2024,^5^ with limited therapeutic options and significant burden of disease. Critical pathogens include carbapenem- and third-generation cephalosporin-resistant Enterobacterales, including *Escherichia coli* and *Klebsiella pneumoniae*, and carbapenem-resistant *Acinetobacter baumannii* (CRAB). Carbapenem-resistant *Pseudomonas aeruginosa* (CRPA) are categorised as high priority. Epidemiological analyses have revealed widespread dissemination and expansion of these priority pathogens.^2,6–8^

Alongside broader public health interventions, this challenge requires continued development and validation of novel treatments and diagnostic assays to improve patient outcomes and strengthen AMR surveillance. Well-characterised panels of isolates are essential to these efforts: standardised, accessible collections enable robust evaluation of new molecules and devices and improve reproducibility across laboratories.^9–12^ Such panels should reflect the contemporary epidemiological landscape and represent strains of greatest public health importance, including globally relevant sequence types (STs) and resistance mechanisms.

However, available Gram-negative panels remain limited, often because of incomplete characterisation, underrepresentation of emerging high-risk clones and WHO priority pathogens, or limited suitability for countermeasure development.^11–18^ Well-characterised, contemporary panels of priority Gram-negative pathogens are needed to support antimicrobial discovery and evaluation.

Here, we present a collection of five isolate panels for key Gram-negative priority pathogens: *E. coli, K. pneumoniae, A. baumannii, P. aeruginosa*, and a mixed panel of other Enterobacterales species, including *Enterobacter hormaechei, Enterobacter kobei, Citrobacter koseri, Citrobacter freundii, Serratia marcescens, Morganella morganii*, and *Proteus mirabilis*. Pathogens selected for inclusion are associated with high-burden clinical indications, including urinary tract infections, respiratory tract infections and bloodstream infections. Selection of strain and AMR phenotype was guided by the WHO BPPL list.^5^ STs are diverse and representative of nationally and internationally relevant clones; ST selection was informed by the wider literature and unpublished data from the UK Health Security Agency (UKHSA) Antimicrobial Resistance and Healthcare-Associated Infections (AMRHAI). The *E. coli, K. pneumoniae* and mixed Enterobacterales panels include third generation cephalosporin- or carbapenem-resistant isolates; isolates within the *A. baumannii* and *P. aeruginosa* are all carbapenem-resistant.

Panels were selected from isolates referred to the UKHSA’s AMRHAI Reference Unit, the UK’s national reference laboratory for investigating unusual AMR in healthcare-associated bacteria, from diagnostic laboratories nationwide. This work was undertaken in collaboration with PACE, an initiative supporting early-stage innovation in medicines and diagnostics to tackle AMR.

We performed genotypic and phenotypic characterisation of antimicrobial susceptibility on all isolates. Panels and accompanying metadata are available for use in developing novel antimicrobial therapeutics and diagnostic assays.

## MATERIALS AND METHODS

### Bacterial isolates and panel selection

Isolates were originally referred to AMRHAI by diagnostic laboratories from across England’s nine regions for investigation of unusual resistance phenotypes and/or antibiotic susceptibility testing for therapeutic guidance, or characterisation to support outbreak investigations. Panels were designed to represent bacterial pathogens on the WHO BPPL 2024 belonging to diverse STs currently circulating in the UK and globally,^5^ and harbouring resistance mechanisms of public health importance, with a focus on mechanisms underpinning resistance to the third-generation cephalosporins, carbapenems, colistin and other ‘last-line’ antibiotics.

### Antimicrobial susceptibility testing

Antibiotic minimum inhibitory concentrations (MICs) were determined via broth microdilution (MICRONAUT, Bruker) against AMRHAI’s standard antibiotic panel,^19^ and interpreted according to the EUCAST clinical breakpoints version 16.0.^20^ Cefiderocol susceptibility testing was performed by disk diffusion with a 30 µg cefiderocol disk (Mast Group). Breakpoints for temocillin, amikacin, and gentamicin, applicable only for infections originating from the urinary tract, were adopted irrespective of original infection site. The breakpoint of 0.5 mg/L for temocillin, applicable only against *E. coli, K. pneumoniae*, and *P. mirabilis*, was adopted for all Enterobacterales species. Where breakpoints were not defined, isolates were categorised as ‘wildtype’ (WT) or ‘non-wildtype’ (non-WT) according to EUCAST Epidemiological Cut-Off (ECOFF) values. At the time of writing, no EUCAST breakpoint or ECOFF value has been defined for tigecycline against *A. baumannii*.

### Whole genome sequencing and analysis

DNA was extracted from RNase-treated lysates using a QIAsymphony DSP DNA Midi Kit (QIAGEN, Hilden, Germany). Sequencing libraries were prepared using Illumina DNA Prep and sequenced with the Illumina NextSeq platform (Illumina, San Diego, USA). Trimming of raw sequencing reads and identification of species and MLST were performed as previously described.^21^ *De novo* genome assembly was performed using *SPAdes (v3*.*15*.*5)*.^22^ Genome assemblies have been deposited in the European Nucleotide Archive at EMBL-EBI under project accession PRJEB95032. Accession numbers for each isolate are listed in Supplementary Tables 2 – 6.

*AMRFinderPlus (v4*.*0*.*3)* was used to identify genotypic AMR determinants.^23^ *In silico* Clermont phylotyping and serotype prediction of *E. coli* genomes were performed with *EZClermont (v1*.*0)* and *ECTyper (v2*.*0*.*0)*, respectively.^24,25^ *In silico* K- and O-antigen typing for *K. pneumoniae* genomes, and K- and OC-antigen genotyping for *A. baumannii*, were performed with *Kaptive (v3*.*2*.*1)*.^26^ Maximum likelihood phylogenetic trees were inferred from core gene SNP alignments to visualise population structure and diversity for *E. coli, K, pneumoniae, CRAB* and *CRPA* panels. Core gene alignments were generated using *Panaroo (v1*.*7*.*0)* with a 95% threshold to determine core genes, then variant sites extracted using *SNP-sites (v2*.*5*.*1)*.^27,28^ Trees were constructed using *IQ-TREE v2*.*4*.*0* with the best supported evolutionary model identified by *IQ-TREE* ModelFinder, constant sites, and ultrafast bootstrapping of 1000 replicates.^29–31^ Core gene SNP alignment lengths and evolutionary models used for each species are listed in Supplementary Table 1.

Data analysis and visualisation was performed using *R Statistical Software v4*.*5*.*3*,^32^ with packages *tidyverse v2*.*0*.*0*,^33^ *patchwork v1*.*3*.*2*,^34^ *gnewscale v0*.*5*.*2*,^35^ *treeio v1*.*34*.*0*,^36^ *ggtree v4*.*0*.*5*,^37^ *ggtreeExtra v1*.*20*.*1*.^38^

## RESULTS AND DISCUSSION

Five bacterial isolate panels were built representing key WHO Gram-negative priority pathogens. Four panels comprise 30 isolates each of *Escherichia coli, Klebsiella pneumoniae*, carbapenem-resistant *Acinetobacter baumannii* and carbapenem-resistant *Pseudomonas aeruginosa*. A fifth isolate panel containing 25 assorted Enterobacterales isolates was assembled, comprising: four *Enterobacter hormaechei*, one *Enterobacter kobei*, two *Citrobacter koseri*, three *Citrobacter freundii*, five *Serratia marcescens*, five *Morganella morganii*, and five *Proteus mirabilis*. All isolates have been accessioned and are made available by the National Collection of Type Cultures (NCTC). Catalogue numbers for each isolate are referenced below where relevant and listed in full in Supplementary Tables 2 – 6.

Panels were designed to include isolates belonging to globally relevant clones harbouring resistance mechanisms and/or exhibiting resistance phenotypes of public health importance. All isolates were originally referred to AMRHAI between 2014 and 2025, with the majority (95/145 isolates, 65.5%) collected since 2023. Samples were originally isolated from a range of sites, including urine and urinary catheter (42/145, 30.0%), faeces and rectal swabs, (29/145, 20.0%), respiratory (21/145, 14.5%), and blood (18/145, 12.4%). Isolation sites are listed in full in Supplementary Tables 2 – 6.

### Escherichia coli panel

The *E. coli* panel includes 14 STs. Five STs, all international high-risk clones, are represented by three or more isolates (ST131 (4/30); ST38 (4/30); ST405 (3/30); ST648 (3/30); ST940 (3/30)). The panel includes specific clones of public health importance such as ST131 carrying *bla*_KPC-2_ (NCTC 15154), ST38 carrying *bla*_OXA-48-like_ (NCTC 15145, *bla*_OXA-48_; NCTC 15148, *bla*_OXA-244_), ST167 carrying *bla*_NDM-5_ (NCTC 15156, 15157), and ST648 carrying *bla*_NDM-5_ (NCTC 15165).^21,39^

Figure 1 describes the phylogeny of the *E. coli* panel alongside key resistance mechanisms and antimicrobial susceptibility. For a full description of all isolates in the *E. coli* panel, see Supplementary Table 2.

**Figure 1.**
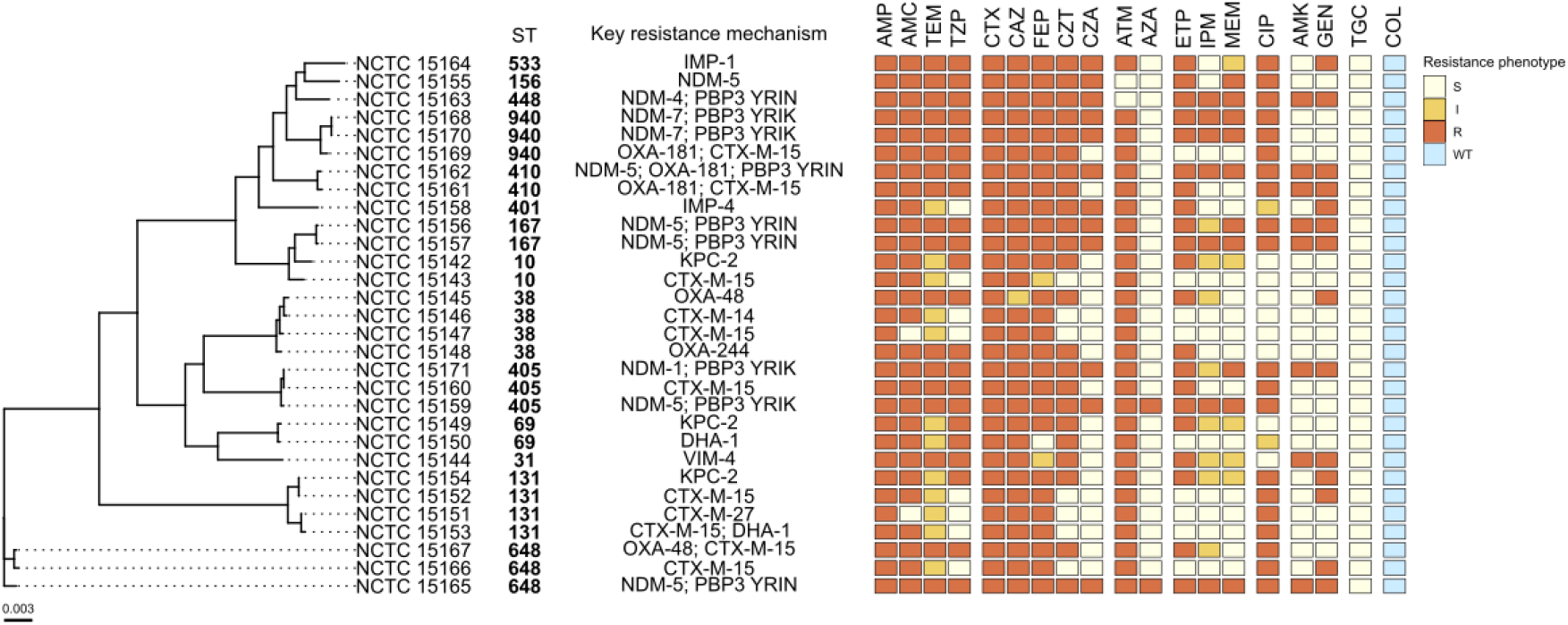
Primary characteristics of the *E. coli* panel. Maximum-likelihood core gene SNP phylogeny and sequence types of the 30 isolates in the *E. coli* panel. Key antimicrobial resistance mechanisms are indicated alongside antimicrobial susceptibility phenotypes to 19 clinically relevant antibiotics. Tree scale bar shows nucleotide substitutions per site. NCTC, National Collection of Type Cultures catalogue number. Resistance phenotype key: S, susceptible; I, susceptible with increased exposure; R, resistant; WT, wildtype. TEM, temocillin; CZT, ceftolozane-tazobactam; CZA, ceftazidime-avibactam; AZA, aztreonam-avibactam; COL, colistin. *Alt text:* Phylogenetic tree of the 30 isolates included in the *E. coli* panel, labelled according to their catalogue number, sequence type and key resistance mechanisms. The isolates’ antimicrobial susceptibility profile to 19 different antimicrobials is also shown in a colour-coded graph.

Carbapenem-resistant Gram-negative bacteria have emerged globally over the past two decades, with most harbouring carbapenemase genes associated with the spread of high-risk clones. Many *E. coli* isolates in the panel (21/30) harbour an acquired carbapenemase gene, often in addition to an extended-spectrum beta-lactamase (ESBL) gene. Across these isolates, three classes of carbapenemase genes are represented, and include the five most commonly found families: class A (*bla*_KPC_; three isolates), class B (*bla*_NDM,_ *bla*_VIM,_ *bla*_IMP_; 13 isolates) and class D (*bla*_OXA-48-like_; six isolates). One isolate (NCTC 15162) carries two carbapenemase genes from different families, *bla*_NDM-5_ + *bla*_OXA-181_. Selection of carbapenemase gene families included in the panel was informed by recent genomic epidemiological analyses of isolates referred to the AMRHAI and review of the published literature.^21^ Eight isolates in the *E. coli* panel have ESBL genes as the major resistance mechanism. Among these, *bla*_CTX-M-15_ is the predominant type (six isolates), with two carrying *bla*_CTX-M-9-like_ ESBL genes. One AmpC-producing isolate (NCTC 15150, *bla*_DHA-1_) is included, representative of the ST69 lineage predominant in urinary tract and bloodstream infections.^40^

Two isolates (NCTC 15159, 15165) harbour an insertion in the PBP3 gene *ftsI*, associated with aztreonam-avibactam resistance. In another two isolates we detected mobile colistin resistance (*mcr*) genes (NCTC 15419, *mcr9*.*2*; NCTC 15158, *mcr9*.*1*), however when tested against colistin, all *E. coli* panel isolates had a MIC ≤1 mg/L, below the ECOFF value.

### Klebsiella pneumoniae panel

The *K. pneumoniae* panel comprises 15 STs, including well-known and emerging international high-risk clones and hypervirulent strains.^21, 41–43^ Four STs are represented by three or more isolates: ST147 (5/30); ST11 (4/30); ST101 (3/30); ST307 (3/30). The panel also includes two ST23 isolates (NCTC 15172, 15175), an emerging lineage increasingly detected across European healthcare settings, carrying hypervirulence and carbapenemase genes.^44^ See Figure 2; Supplementary Table 3.

**Figure 2.**
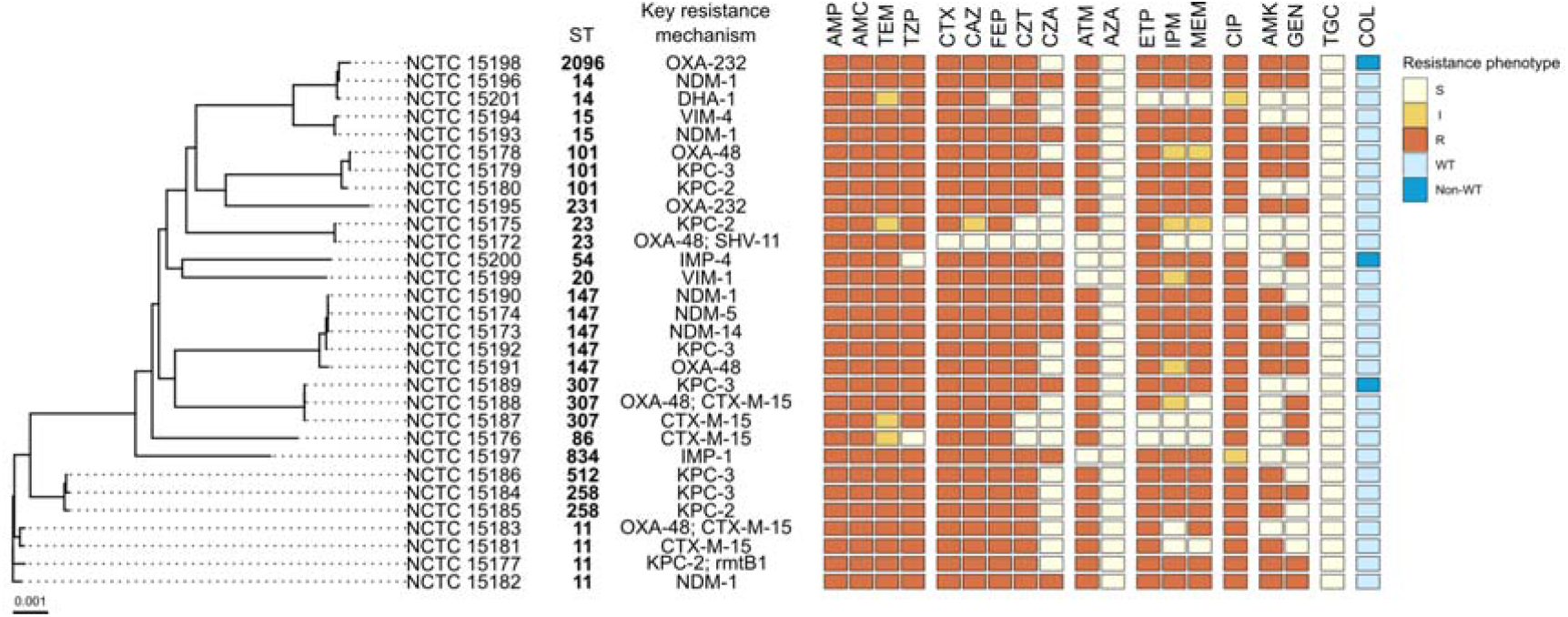
Primary characteristics of the *K. pneumoniae* panel. Maximum-likelihood core gene SNP phylogeny and sequence types of the 30 isolates in the *K. pneumoniae* panel. Key resistance mechanisms are indicated alongside antimicrobial susceptibility phenotypes to 19 clinically relevant antibiotics. Tree scale bar shows nucleotide substitutions per site. NCTC, National Collection of Type Cultures catalogue number. Resistance phenotype key: S, susceptible; I, susceptible with increased exposure; R, resistant; WT, wildtype. TEM, temocillin; CZT, ceftolozane-tazobactam; CZA, ceftazidime-avibactam; AZA, aztreonam-avibactam; COL, colistin. *Alt text:* Phylogenetic tree of the 30 isolates included in the *K. pneumoniae* panel, labelled according to their catalogue number, sequence type and key resistance mechanisms. The isolates’ antimicrobial susceptibility profile to 19 different antimicrobials is also shown in a colour-coded graph.

Twenty-nine carbapenemase genes were identified across 26 carbapenemase producing isolates, with the selection guided by knowledge of currently circulating clones and review of the published literature.^21^ *bla*_KPC_ and *bla*_OXA-48-like_ were each found in nine isolates, and class B metallo-β-lactamase (MBL)-type carbapenemase genes are present in 11 isolates. These are mostly *bla*_NDM_ variants (seven isolates), but *bla*_IMP_ (two isolates) and *bla*_VIM_ (two isolates) are also represented. Three isolates were found to carry two carbapenemase genes (NCTC 15174, *bla*_NDM-5_ + *bla*_OXA-232_; NCTC 15180, *bla*_NDM-1_ + *bla*_KPC-2_; NCTC 15196, *bla*_NDM-1_ + *bla*_OXA-181_).

The panel includes one isolate harbouring the plasmid-mediated AmpC *bla*_DHA-1_ (NCTC 15201). Three remaining isolates carrying *bla*_CTX-M-15_ were selected as representative ESBL carriers.

Many isolates were resistant to aminoglycosides (21/30), and in nine isolates an acquired 16S rRNA methyltransferase gene associated with high-level resistance to all clinically relevant aminoglycosides, such as *armA* and *rmtF1*, was identified. Three isolates (NCTC 15189, 15198, 15200) exhibited a colistin MIC ≥ 8 mg/L and were categorised as non-WT; however, no previously described resistance mechanism was detected.

Diverse K- and O-antigen types are represented among the *K. pneumoniae* panel isolates. Knowledge of antigen types can aid research into putative targets for immunotherapeutics and vaccines, while K-types are likely to be important in determining phage susceptibility in many cases.^45,46^

### Carbapenem-resistant Acinetobacter baumannii (CRAB) panel

The CRAB panel comprises 14 STs, including ST2 (10/30), the dominant *A. baumannii* clone globally,^47^ and ST164 (3/30) and ST32 (3/30), both emerging, highly resistant clones linked to MDR hospital outbreaks.^48–50^ See Figure 3; Supplementary Table 4.

**Figure 3.**
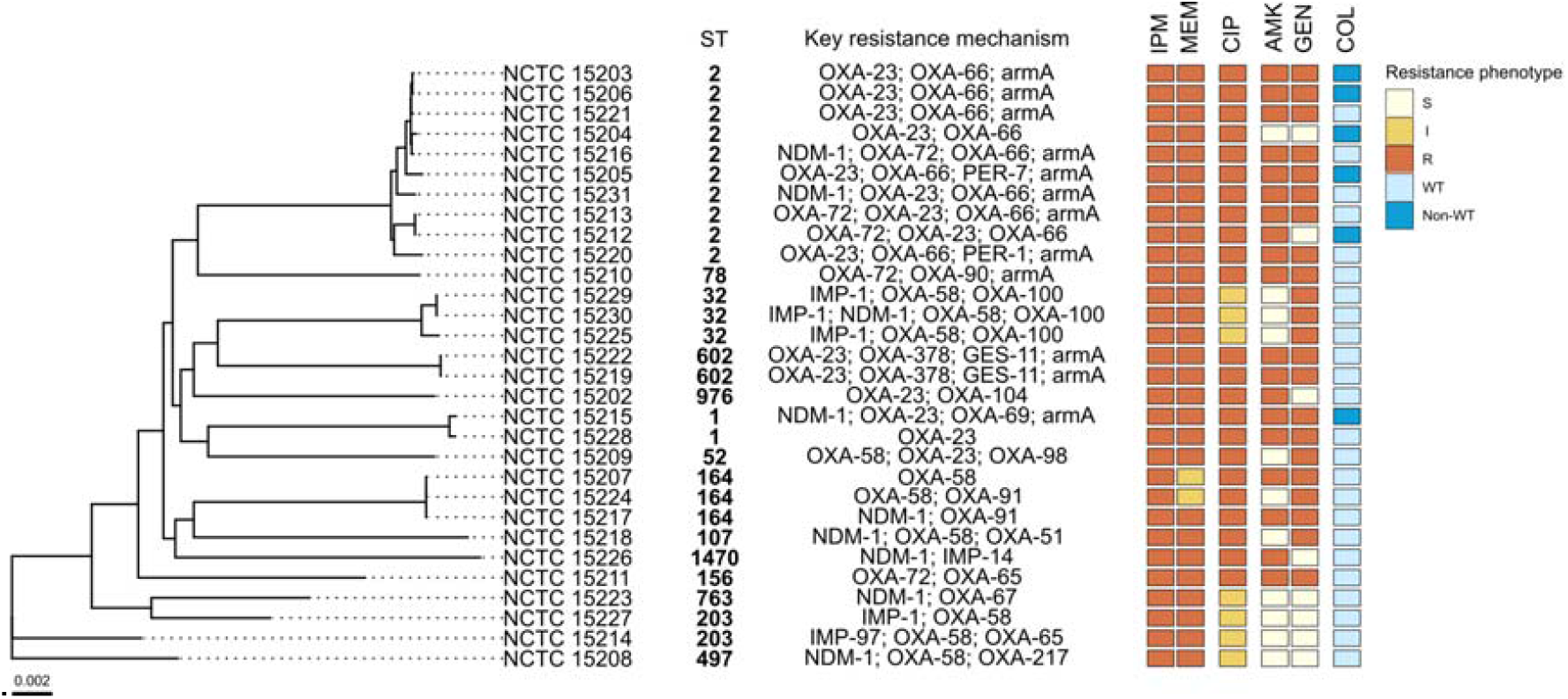
Primary characteristics of the *A. baumannii* panel. Maximum-likelihood core gene SNP phylogeny and sequence types of the 30 isolates in the *A. baumannii* panel. Key resistance mechanisms are indicated alongside antimicrobial susceptibility phenotypes to 6 clinically relevant antibiotics. Tree scale bar shows nucleotide substitutions per site. NCTC, National Collection of Type Cultures catalogue number. Resistance phenotype key: S, susceptible; I, susceptible with increased exposure; R, resistant; WT, wildtype. COL, colistin. *Alt text:* Phylogenetic tree of the 30 isolates included in the *A. baumannii* panel, labelled according to their catalogue number, sequence type and key resistance mechanisms. The isolates’ antimicrobial susceptibility profile to 6 different antimicrobials is also shown in a colour-coded graph.

Forty-five acquired carbapenemase genes were identified across the 30 isolates, representing five gene families: *bla*_OXA-23-like_ (15 isolates), *bla*_OXA-58-like_ (10 isolates), *bla*_OXA-24-like_ (five isolates), *bla*_NDM_ (nine isolates), and *bla*_IMP_ (six isolates). Thirteen isolates carry two acquired carbapenemase genes; one isolate (NCTC 15230) carries three (*bla*_OXA-58_ + *bla*_NDM-1_ + *bla*_IMP-1_). The panel also includes isolates that carry an ESBL gene in addition to an acquired carbapenemase gene: *bla*_PER_ (NCTC 15205, 15220) and *bla*_GES_ (NCTC 15219, 15222).

Six of the 30 *A. baumannii* isolates were defined as non-WT against colistin but a previously described resistance mechanism was only identified in two of these: NCTC 15203 and NCTC 15206 both carry a substitution mutation in *pmrC*, a gene encoding a phosphoethanolamine transferase.^51^ The panel includes four isolates with resistance to cefiderocol (NCTC 15205, 15206, 15219, 15220).

### Carbapenem-resistant Pseudomonas aeruginosa (CRPA) panel

The CRPA panel is highly diverse with isolates representing 23 STs, including ST235 (4/30), one of the most prevalent MDR *P. aeruginosa* clones globally,^52^ and high-risk clones ST357 (4/30) and ST654 (2/30). See Figure 4, Supplementary Table 5.

**Figure 4.**
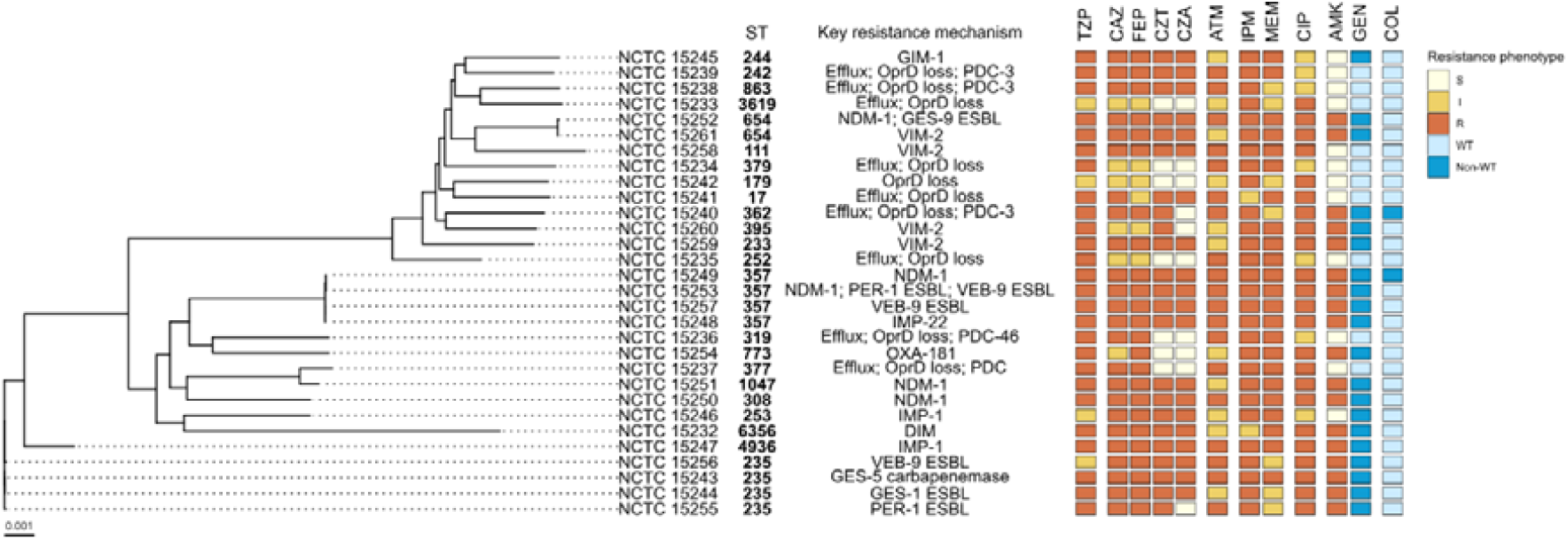
Primary characteristics of the *P. aeruginosa* panel. Maximum-likelihood core gene SNP phylogeny and sequence types of the 30 isolates in the *P. aeruginosa* panel. Key resistance mechanisms are indicated alongside antimicrobial susceptibility phenotypes to 12 clinically relevant antibiotics. Tree scale bar shows nucleotide substitutions per site. NCTC, National Collection of Type Cultures catalogue number. Resistance phenotype key: S, susceptible; I, susceptible with increased exposure; R, resistant; WT, wildtype. CZT, ceftolozane-tazobactam; CZA, ceftazidime-avibactam; COL, colistin. *Alt text:* Phylogenetic tree of the 30 isolates included in the *P. aeruginosa* panel, labelled according to their catalogue number, sequence type and key resistance mechanisms. The isolates’ antimicrobial susceptibility profile to 12 different antimicrobials is also shown in a colour-coded graph.

The panel includes 16 isolates with an acquired carbapenemase gene, representing known international high-risk clones.^53^ Among carbapenemase gene carriers, one isolate harbours *bla*_GES-5_, one harbours *bla*_OXA-181_, and 14 isolates carry an MBL-type carbapenemase: *bla*_NDM-1_ (five isolates), *bla*_VIM-2_ (four isolates), *bla*_IMP-1_ (two isolates), *bla*_IMP-22_ (one isolate), as well as less common variants *bla*_DIM-1_ and *bla*_GIM-1_ (one isolate each).

The remaining 14 CRPA isolates were selected based on antibiotic susceptibility profiles consistent with derepressed AmpC activity, increased expression of efflux pumps and/or decreased expression of outer membrane porin OprD. In many of these isolates, known mutations in *oprD* and *ampR* (a regulator of *ampC*) were identified through genomic analysis. The panel also includes several representative ESBL-positive isolates, including one isolate carrying *bla*_GES-1_ (NCTC 15244), one carrying *bla*_PER-1_ (NCTC 15255), and two carrying *bla*_VEB-9_ (NCTC 15256, 15257).

Colistin MICs were above the ECOFF value for two isolates, categorised as non-WT (NCTC 15240, 15249). Eighteen isolates were resistant to amikacin; the ribosomal RNA methyltransferase gene *rmtF*, which confers high-level resistance to amikacin and other aminoglycoside antibiotics,^54^ was identified in four isolates, all of which are also MBL carriers.

### Mixed Enterobacterales panel

Within the mixed Enterobacterales panel, 15 STs are represented, including clinically predominant *C. freundii* type ST22 and high-risk *P. mirabilis* clone ST135 associated with extensive MDR.^55,56^ Two *Citrobacter koseri* isolates could not be typed: NCTC 15287 could not be assigned to a defined ST at the time of analysis (July 2026) and represents a novel ST; an ST could not be assigned to NCTC 15286 due to incomplete locus coverage. See Figure 5, Supplementary Table 6.

**Figure 5.**
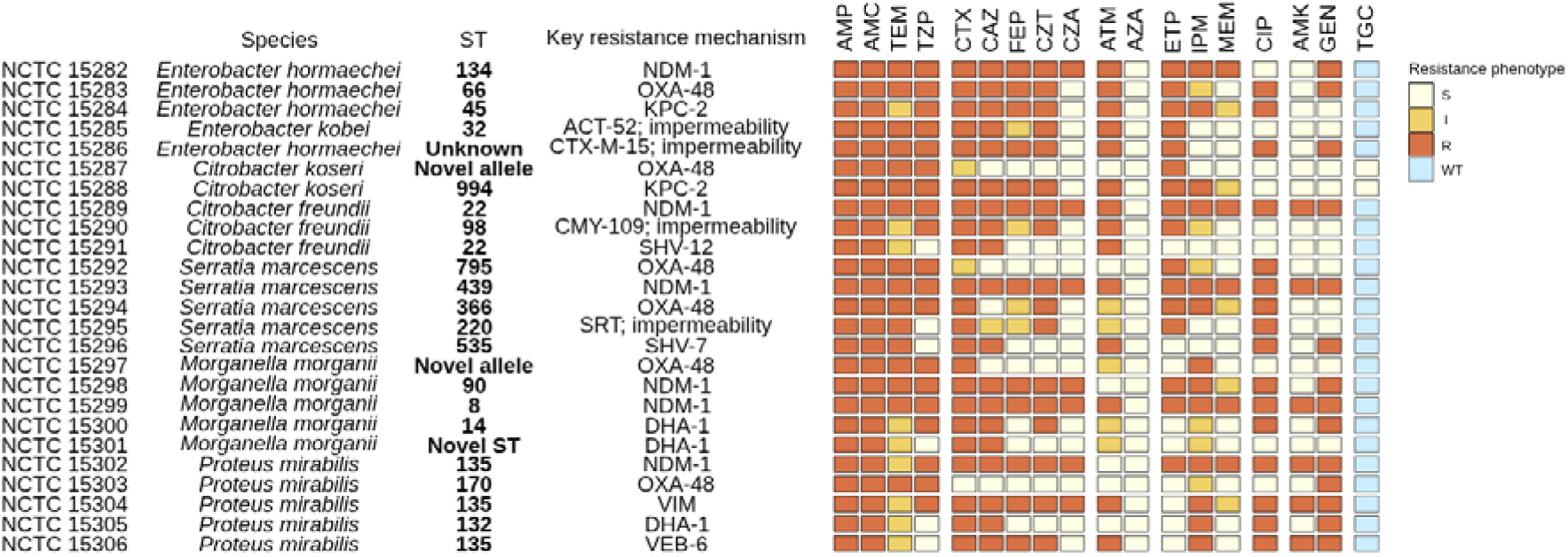
Primary characteristics of the mixed Enterobacterales panel. Species and sequence types of the 25 isolates in the mixed Enterobacterales panel. Key resistance mechanisms are indicated alongside antimicrobial susceptibility phenotypes to 18 clinically relevant antibiotics. EUCAST breakpoint for urinary tract infections adopted for all species irrespective of infection site (see Materials and methods). NCTC, National Collection of Type Cultures catalogue number. Resistance phenotype key: S, susceptible; I, susceptible with increased exposure; R, resistant; WT, wildtype. TEM, temocillin; CZT, ceftolozane-tazobactam; CZA, ceftazidime-avibactam; AZA, aztreonam-avibactam. *Alt text:* The 25 isolates included in the mixed Enterobacterales panel, labelled according to their catalogue number, sequence type and key resistance mechanisms. The isolates’ antimicrobial susceptibility profile to 18 different antimicrobials is also shown in a colour-coded graph.

Fifteen isolates harbour a carbapenemase gene, representing four gene families: *bla*_NDM-1_ (six isolates), *bla*_OXA-48_ (six isolates), *bla*_KPC-2_ (two isolates), and *bla*_VIM_ (one isolate; variant undetermined). Four isolates were selected based on ESBL gene carriage: an *E. hormaechei* isolate with *bla*_CTX-M-15_ (NCTC 15286), a *C. freundii* isolate carrying *bla*_SHV-12_ (NCTC 15291), a *S. marcescens* isolate carrying *bla*_SHV-7_ (NCTC 15296) and a *P. mirabilis* isolate with *bla*_VEB-6_ (NCTC 15306).

Two *Enterobacter* isolates carry *mcr-9* variants (NCTC 15284, *mcr9-2*; NCTC 15285, *mcr9-1*); unlike other *mcr* variants, *mcr-9* does not typically confer colistin resistance. NCTC 15285 also carries known colistin resistance determinant *qseB/qseC*. However, colistin MICs remained low for both (≤0.5 mg/L). The isolates selected for inclusion in the mixed Enterobacterales panel are opportunistic pathogens often associated with high-burden indications, including urinary tract and bloodstream infections, and with hospital-acquired infections, particularly in vulnerable or immunocompromised patients. Despite representing a significant and growing clinical challenge, these species are generally less well understood than those represented in the other panels. The extensive WGS and antimicrobial susceptibility data provided in this mixed panel could support the development of therapeutics and diagnostics for indications caused by Enterobacterales pathogens, as well as the evaluation of new products against less common strains. This broader strain coverage may help to inform clinical positioning and potential differentiation opportunities of novel therapeutics and diagnostics.

## CONCLUDING PARAGRAPHS

MDR Gram-negative bacteria are critical threats to public health.^5^ Around 40% of *E. coli* and 55% of *K. pneumoniae* are now resistant to third-generation cephalosporins, the first-choice treatment for severe bloodstream infections.^57^ Carbapenem resistance is becoming more frequent, narrowing treatment options and forcing reliance on other last-resort antibiotics.^57^

Testing against diverse, contemporary clinical isolates is essential for the development, evaluation and validation of new antimicrobial diagnostics and therapeutics that could counteract these challenges. There is, however, a lack of well-characterised, clinically relevant MDR pathogen panels, leaving significant gaps in the supporting evidence of novel innovations. UKHSA’s AMRHAI Reference Unit’s national genomic, molecular and phenotypic surveillance reference services provided a unique opportunity to access strains with diverse AMR phenotypes and curate panels of isolates highly representative of those currently in circulation.

In this paper, we described five screening panels each containing 25 to 30 isolates. Isolates have been extensively characterised both genotypically, through WGS, and phenotypically, through antibiotic susceptibility testing using ‘gold-standard’ methodology. Importantly, these panels offer a highly curated collection of contemporary and globally disseminated clones with resistance mechanisms that pose the greatest risk to public health, complementing existing panels selected for maximum genetic diversity.^13–15^

The isolates have been deposited in the National Collection of Type Cultures and are accessible to the wider global scientific and innovation community as a resource for evaluating potential new antimicrobials and diagnostics for clinically important resistant pathogens.

## FIGURES

Please see separate doc

## ACKNOWLEDGEMENTS

The authors thank members of the wider team for their contribution to the work: Neill Gingles, Medicines Discovery Catapult; Phil Packer, Innovate UK; Clive Mason, LifeArc. The authors would also like to thank Emily C Farthing for her valuable editorial support.

This project is the result of a partnership between the UKHSA and PACE (Pathways to Antimicrobial Clinical Efficacy). PACE is a collaboration between LifeArc, Medicines Discovery Catapult and Innovate UK supporting early-stage innovation in medicines and diagnostics to tackle AMR and save lives.

Opinions expressed are those of the authors and not necessarily those of UKHSA or the Department of Health and Social Care.

## TRANSPARENCY DECLARATIONS

SM and KLH are employees of the UKHSA’s Antimicrobial Resistance and Healthcare Associated Infections Reference Unit, which has received financial support for conference attendance, research projects or contracted evaluations from numerous sources, including Advanz Pharma Services (UK) Ltd, A. Menarini Farmaceutica Internazionale SRL, GlaxoSmithKline Services Ltd, Merck Sharpe & Dohme Corp, Pfizer Ltd, Paion Pharma GmbH, and Shionogi & Co. Ltd. All other authors declare no conflict of interest. Editorial support was provided by Emily C Farthing, independent science writer and editor.

## AUTHOR CONTRIBUTIONS

JKS: drafting of the manuscript, acquisition and analysis of data; SM, CH: conception, design, data analysis, manuscript review; RA, HMc ES, EH: data acquisition and analysis; HM, EB, LD: conception and design, manuscript review; PJC, BI, JMS: conception and design, critical review and final approval of the manuscript; KLH: conception and design, data analysis, critical review and final approval of the manuscript.

## SUPPLEMENTARY DATA

Supplementary tables 1 – 6 are available as Supplementary data.

## DATA AVAILABILITY

Genome assemblies have been deposited in the European Nucleotide Archive at EMBL-EBI under project accession PRJEB95032. Accession numbers for each strain are listed in Supplementary Table 2 – 6.

## Supplementary tables headers

**Supplementary table 1**

Core SNP alignment lengths and evolutionary models used to infer phylogenetic trees.

**Supplementary table 2**

Primary characteristics of 30 isolates found in the *E. coli* panel.

**Supplementary table 3**

Primary characteristics of 30 isolates found in the *K. pneumoniae* panel.

**Supplementary table 4**

Primary characteristics of 30 isolates found in the *A. baumannii* panel.

**Supplementary table 5**

Primary characteristics of 30 isolates found in the *P. aeruginosa* panel.

**Supplementary table 6**

Primary characteristics of 30 isolates found in the mixed Enterobacterales panel.

Classification: Internal

## REFERENCES

1. Naghavi M, Vollset SE, Ikuta KS et al. Global burden of bacterial antimicrobial resistance 1990–2021: a systematic analysis with forecasts to 2050. Lancet 2024;404:1199–226.

2. UK Health Security Agency (UKHSA). English Surveillance Programme for Antimicrobial Utilisation and Resistance (ESPAUR) Report 2024 to 2025. London, 2025.

3. Maher C, Hassan K. The Gram-negative permeability barrier: tipping the balance of the in and the out. MBio 2023;14:e01205–23.

4. Melchiorri D, Rocke T, Alm RA et al. Addressing urgent priorities in antibiotic development: insights from WHO 2023 antibacterial clinical pipeline analyses. Lancet Microbe 2025;6:100992.

5. Sati H, Carrara E, Savoldi A et al. The WHO Bacterial Priority Pathogens List 2024: a prioritisation study to guide research, development, and public health strategies against antimicrobial resistance. Lancet Infect Dis 2025;25:1033–43.

6. Reynolds R, Mushtaq S, Hope R et al. Antimicrobial resistance among Gram-negative agents of bacteraemia in the UK and Ireland: trends from 2001 to 2019. J Antimicrob Chemother 2025;80:iv36–48.

7. Otu A, McCormick J, Henderson KL et al. Understanding the landscape of carbapenemase-producing organisms (CPOs), and spotlighting opportunities for control in England. Infect Prev Pract 2025;7:100480.

8. Wise MG, Karlowsky JA, Mohamed N et al. Global trends in carbapenem- and difficult-to-treat-resistance among World Health Organization priority bacterial pathogens: ATLAS surveillance program 2018–2022. J Glob Antimicrob Resist 2024;37:168–75.

9. Livermore DM, Warner M, Mushtaq S. Activity of MK-7655 combined with imipenem against Enterobacteriaceae and Pseudomonas aeruginosa. J Antimicrob Chemother 2013;68:2286–90.

10. Mushtaq S, Sadouki Z, Vickers A et al. In Vitro Activity of cefiderocol, a siderophore cephalosporin, against multidrug-resistant Gram-negative bacteria. Antimicrob Agents Chemother 2020;64.

11. Deckers C, Soleimani R, Denis O et al. Multicentre interlaboratory analysis of routine susceptibility testing with a challenge panel of resistant strains. J Glob Antimicrob Resist 2022;28:125–9.

12. Desmet S, Verhaegen J, Glupzcynski Y et al. Development of a national EUCAST challenge panel for antimicrobial susceptibility testing. Clin Microbiol Infect 2016;22:704–10.

13. Martin MJ, Stribling W, Ong AC et al. A panel of diverse Klebsiella pneumoniae clinical isolates for research and development. Microb Genom 2023;9:mgen000967.

14. Lebreton F, Snesrud E, Hall L et al. A panel of diverse Pseudomonas aeruginosa clinical isolates for research and development. JAC Antimicrob Resist 2021;3:dlab179.

15. Galac MR, Snesrud E, Lebreton F et al. A diverse panel of clinical Acinetobacter baumannii for research and development. Antimicrob Agents Chemother 2020;64:aac.00840-20.

16. Soyza A De, Hall AJ, Mahenthiralingam E et al. Developing an international Pseudomonas aeruginosa reference panel. Microbiologyopen 2013;2:1010–23.

17. D’Souza R, Pinto NA, Hwang I et al. Panel strain of Klebsiella pneumoniae for beta-lactam antibiotic evaluation: their phenotypic and genotypic characterization. PeerJ 2017;5:e2896.

18. D’Souza R, Pinto NA, Phuong N Le et al. Phenotypic and genotypic characterization of Acinetobacter spp. panel strains: a cornerstone to facilitate antimicrobial development. Front Microbiol 2019;10:559.

19. UK Health Security Agency. Bacteriology Reference Department (BRD) user manual. 2026. https://assets.publishing.service.gov.uk/media/6a564a6f2f6185941a9a64b4/bacteriology-reference-department-user-manual-2026-July.pdf.

20. European Committee on Antimicrobial Susceptibility Testing. Breakpoint Tables for Interpretation of MICs and Zone Diameters. Version 16.0. 2026. https://www.eucast.org.

21. Hopkins KL, Ellaby N, Ellington MJ et al. Diversity of carbapenemase-producing Enterobacterales in England as revealed by whole-genome sequencing of isolates referred to a national reference laboratory over a 30-month period. J Med Microbiol 2022;71.

22. Prjibelski A, Antipov D, Meleshko D et al. Using SPAdes De Novo Assembler. Curr Protoc Bioinformatics 2020;70:e102.

23. Feldgarden M, Brover V, Gonzalez-Escalona N et al. AMRFinderPlus and the Reference Gene Catalog facilitate examination of the genomic links among antimicrobial resistance, stress response, and virulence. Sci Rep 2021;11:12728.

24. Waters NR, Abram F, Brennan F et al. Easy phylotyping of Escherichia coli via the EzClermont web app and command-line tool. Access Microbiol 2020;2:acmi000143.

25. Bessonov K, Laing C, Robertson J et al. ECTyper: in silico Escherichia coli serotype and species prediction from raw and assembled whole-genome sequence data. Microb Genom 2021;7:000728.

26. Stanton TD, Hetland MAK, Löhr IH et al. Fast and accurate in silico antigen typing with Kaptive 3. Microb Genom 2025;11:001428.

27. Tonkin-Hill G, MacAlasdair N, Ruis C et al. Producing polished prokaryotic pangenomes with the Panaroo pipeline. Genome Biol 2020;21:180.

28. Page AJ, Taylor B, Delaney AJ et al. SNP-sites: rapid efficient extraction of SNPs from multi-FASTA alignments. Microb Genom 2016;2:e000056.

29. Minh BQ, Schmidt HA, Chernomor O et al. IQ-TREE 2: New models and efficient methods for phylogenetic inference in the genomic era. Mol Biol Evol 2020;37:1530–4.

30. Kalyaanamoorthy S, Minh BQ, Wong TKF et al. ModelFinder: fast model selection for accurate phylogenetic estimates. Nat Methods 2017;14:587–9.

31. Hoang DT, Chernomor O, Haeseler A von et al. UFBoot2: improving the ultrafast bootstrap approximation. Mol Biol Evol 2018;35:518–22.

32. R Core Team. R: A language and environment for statistical computing. Preprint, R Foundation for Statistical Computing, 2026. 10.32614/R.manuals.

33. Wickham H, Averick M, Bryan J et al. Welcome to the Tidyverse. J Open Source Softw 2019;4:1686.

34. Pedersen T. patchwork: The Composer of Plots, R package version 1.0.0. Preprint, 2019. 10.32614/CRAN.package.patchwork.

35. Campitelli E. ggnewscale: multiple fill and colour scales in ‘ggplot2’, R package version 0.5.2. Preprint, 2019. 10.5281/zenodo.2543762.

36. Wang LG, Lam TTY, Xu S et al. Treeio: An R Package for Phylogenetic Tree Input and Output with Richly Annotated and Associated Data. Mol Biol Evol 2020;37:599–603.

37. Yu G, Smith DK, Zhu H et al. ggtree : an r package for visualization and annotation of phylogenetic trees with their covariates and other associated data. Methods Ecol Evol 2017;8:28–36.

38. Xu S, Dai Z, Guo P et al. ggtreeExtra: compact visualization of richly annotated phylogenetic data. Mol Biol Evol 2021;38:4039–42.

39. Shafiq M, Zeng M, Permana B et al. Coexistence of blaNDM–5 and tet(X4) in international high-risk Escherichia coli clone ST648 of human origin in China. Front Microbiol 2022;13:1031688.

40. Doumith M, Day M, Ciesielczuk H et al. Rapid identification of major Escherichia coli sequence types causing urinary tract and bloodstream infections. J Clin Microbiol 2015;53:160–6.

41. Turton JF, Perry C, McGowan K et al. Klebsiella pneumoniae sequence type 147: a high-risk clone increasingly associated with plasmids carrying both resistance and virulence elements. J Med Microbiol 2024;73:001823.

42. Lam MMC, Wick RR, Watts SC et al. A genomic surveillance framework and genotyping tool for Klebsiella pneumoniae and its related species complex. Nat Comm 2021;12:4188.

43. Zhang R, Liu L, Zhou H et al. Nationwide surveillance of clinical carbapenem-resistant Enterobacteriaceae (CRE) strains in China. EBioMedicine 2017;19:98–106.

44. European Centre for Disease Prevention and Control. Emergence of Hypervirulent Klebsiella Pneumoniae ST23 Carrying Carbapenemase Genes in EU/EEA Countries, First Update. Stockholm: ECDC, 2024.

45. Wantuch PL, Knoot CJ, Robinson LS et al. Heptavalent O-antigen bioconjugate vaccine exhibiting differential functional antibody responses against diverse Klebsiella pneumoniae Isolates. J Infect Dis 2024;230:578–89.

46. Rothschild-Rodriguez D, Lambon KS, Kushwaha SK et al. KlebPhaCol: a community-driven resource for Klebsiella research identified a novel phage family. Nucleic Acids Res 2025;53:gkaf1122.

47. Hamidian M, Nigro SJ. Emergence, molecular mechanisms and global spread of carbapenem-resistant Acinetobacter baumannii. Microb Genom 2019;5:e000306.

48. Wang X, Xu T, Zhang Y et al. Comparative genomic and resistance characterization of ST2 and ST164 carbapenem-resistant Acinetobacter baumannii from hospital environments and clinical specimens. Microb Genom 2026;12:001679.

49. Liu H, Moran RA, Doughty EL et al. Longitudinal genomics reveals carbapenem-resistant Acinetobacter baumannii population changes with emergence of highly resistant ST164 clone. Nat Comm 2024;15:9483.

50. Tobin LA, Abu Sabah E, Lebreton F et al. Genomic analysis of early ST32 Acinetobacter baumannii strains recovered in US military treatment facilities reveals distinct lineages and links to the origins of the Tn 6168 ampC transposon. J Antimicrob Chemother 2025;80(3):666–75.

51. Gerson S, Lucaßen K, Wille J et al. Diversity of amino acid substitutions in PmrCAB associated with colistin resistance in clinical isolates of Acinetobacter baumannii. Int J Antimicrob Agents 2020;55:105862.

52. Treepong P, Kos VN, Guyeux C et al. Global emergence of the widespread Pseudomonas aeruginosa ST235 clone. Clin Microbiol Infect 2018;24:258–66.

53. Wright LL, Turton JF, Livermore DM et al. Dominance of international ‘high-risk clones’ among metallo--lactamase-producing Pseudomonas aeruginosa in the UK. J Antimicrob Chemother 2015;70:103–10.

54. Wang S, Wei L, Gao Y et al. Novel amikacin resistance genes identified from human gut microbiota by functional metagenomics. J Appl Microbiol 2022;133(2):898–907.

55. Wang Q, Zhou L, Chen X et al. Global emergence and transmission dynamics of carbapenemase-producing Citrobacter freundii sequence type 22 high-risk international clone: a retrospective, genomic, epidemiological study. Lancet Microbe 2025;6:101149.

56. Deka N, Brauer AL, Connerton K et al. Pangenome analysis of Proteus mirabilis reveals lineage-specific antimicrobial resistance profiles and discordant genotype-phenotype correlations. Antimicrob Agents Chemother 2026;70:e0176825.

57. World Health Organization. Global Antibiotic Resistance Surveillance Report 2025 WHO Global Antimicrobial Resistance and Use Surveillance System (GLASS). Geneva: World Health Organization, 2025.

